# pydca v1.0: a comprehensive software for Direct Coupling Analysis of RNA and Protein Sequences

**DOI:** 10.1101/805523

**Authors:** Mehari B. Zerihun, Fabrizio Pucci, Emanuel Karl Peter, Alexander Schug

## Abstract

The ongoing advances in sequencing technologies have provided a massive increase in the availability of sequence data. This made it possible to study the patterns of correlated substitution between residues in families of homologous proteins or RNAs and to retrieve structural and stability information. Direct coupling Analysis (DCA) infers coevolutionary couplings between pairs of residues indicating their spatial proximity, making such information a valuable input for subsequent structure prediction. Here we present **pydca**, a standalone Python-based software package for the DCA of protein- and RNA-homologous families. It is based on two popular inverse statistical approaches, namely, the mean-field and the pseudo-likelihood maximization and is equipped with a series of functionalities that range from multiple sequence alignment trimming to contact map visualization. Thanks to its efficient implementation, features and user-friendly command line interface, **pydca** is a modular and easy-to-use tool that can be used by researchers with a wide range of backgrounds.

**Availability:** https://github.com/KIT-MBS/pydca

## Introduction

The exponential increase of sequence information due to the recent advances in sequencing technologies have triggered huge interest in statistical methods based on coevolution to retrieve structural and energetic information from families of homologous proteins and RNA. Direct coupling analysis (DCA) represents an accurate way to determine these biomolecular properties starting from a multiple sequence alignment (MSA) of proteins or RNAs. As a key element of DCA, coevolving mutations as part of the evolutionary history can represent a physical coupling and spatial proximity of residue pairs. Inverse methods drawn from statistical physics are used to infer pairwise interactions resulting from a physical contact/spatial proximity. In contrast to prior models based on Mutual Information, they aim at transitive correlation effects. A wide variety of algorithms has been implemented such as the message-passing DCA (mpDCA) [1], mean-field DCA (mfDCA) [2], pseudo-likelihood maximization DCA (plmDCA) [3] or Boltzmann machine learning [4]. Top scoring pairs obtained from DCA have been successfully used as constraints in molecular modeling tools to predict protein [4, 5, 6, 7, 8, 9, 10] and RNA three-dimensional structures [11, 12]. In addition, DCA-based methods have been also employed to study effects of mutations on biomolecular properties. Here we present **pydca**, a Python-based standalone tool that implements a mean-field and a pseudo-likelihood approaches to DCA. The availability of this software, its user-friendly command line and its computational efficiency, together with a rapidly growing number of sequences in databases such as the protein-(Pfam) and the RNA-family database (Rfam) enable scientists from a wide range of backgrounds to carry out DCA in a fast and comprehensive way.

The software is implemented using the Python programming language and its structure follows a modular architecture composed of sub-packages, where each encapsulates a specific task. The dependencies such as Biopython [13] are Python-based and commonly available, which simplifies the user-friendly installation and application in the contact prediction. The mean-field and the pseudo likelihood inverse statistic parameter estimations are performed in two specific modules so that the DCA computation of both protein- and RNA sequences is possible. The computational utilities provided by **pydca** are:

- Curation and trimming of the MSA input data
- Mean-field computation of DCA scores summarized by direct information, Frobenius norm or their average product corrected forms
- Pseudo-likelihood maximization computation of DCA scores summarized by direct information, Frobenius norm or their average product corrected forms.
- Mapping the residues of a reference sequence to the corresponding columns in the MSA when an optional reference sequence is supplied
- Computation of the energy function of the global probability model
- Contact map comparison of DCA-predicted residue pairs with an existing PDB contact map
- Visualization of the true positive rates per rank of residue pairs ranked by coevolutionary scores

The software provides a convenient command line interface for executing specific computations at a time. Each (sub)command is documented and can be looked up through help messages. A typical DCA computation using the mean-field algorithm after the installation is for example: `mfdca compute di <biomolecule> <alignment file> --verbose`, where `<biomolecule>` takes PROTEIN or RNA (case insensitive) and `<alignment file>` is a FASTA formatted file containing MSA data. This triggers the mean field computation of DCA scores summarized by direct information [2] score. When executed in verbose mode (optional argument `--verbose`), it provides extensive logging messages and thereby enables the user to keep track of the computation process. Detailed examples of both mfDCA and plmDCA computation using **pydca** can be found in the supplementary material.

Our implementation of DCA using the mean-field and the pseudo likelihood inverse statistics algorithms provides a suitable lightweight and easy-to-use software that will facilitate extensive computations on the growing sequence data. The mean-field algorithm of pydca is computationally very efficient. On the other hand, the pseudolikelihood implementation typically results in better performance in terms of accuracy.

Finally **pydca** provides in a single package a series of additional DCA computation utilities for protein and RNA sequences such as the trimming of the MSA and the TPP rate computation that will facilitate the user in the computation and in the analysis of DCA results.

## Background: Direct Coupling Analysis

Direct Coupling Analysis (DCA) infers coevolutionary related residues pairs from a multiple sequence alignment (MSA) for proteins and RNA. High coevolutionary scores indicate spatial proximity/physical contacts of the involved residues, the contact information improves modeling of three-dimensional biomolecular structures in structure prediction tools. Mathematically, the statistical model of DCA assigns probabilities to sequences in a given family of homologous proteins or RNAs. The probability *P* (*S*) that a biological sequence *S* = *a*_1_*a*_2_…*a*_*L*_ of length *L* is sampled through the course of evolution is given by the

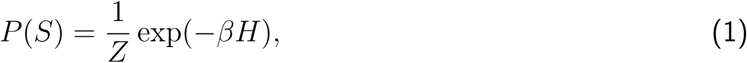

where *β* is the inverse temperature, *Z* is the normalization constant (also known as partition function) and *H* stands for the energy function of the system, which is written as

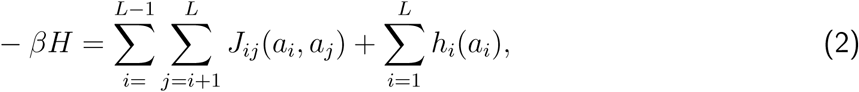

as function of single- and 2-body couplings *h*_*i*_(*a*_*i*_), *J*_*ij*_(*a*_*i*_, *a*_*j*_) that quantify the coupling strength between pairs of sites *i* and *j* for positions *a*_*i*_ and *a*_*j*_, respectively. The single site fields are a measure of the local field strength at a site *i* to a residue or a gap state *a*_*i*_. With the energy function in equation 2, the partition function is given by

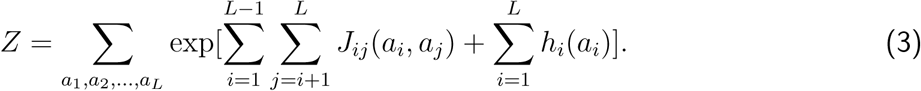

For a sequence of length *L*, a total number of 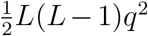 couplings and *Lq* fields is determined, where *q* is the total number of states at a site (5 for RNA and 21 for proteins). The total number of unique parameters that can be obtained equals 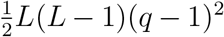 and *L*(*q* − 1) for the couplings and fields, respectively [1].

Computing the summation in equation 3 is computationally costly. It scales as O(*q*^*L*^). As a result inverse statistical inference methods to estimate the fields and couplings rely on approximation methods. The couplings and fields are approximated using inverse statistical algorithms such as the message passing direct-coupling analysis (mpDCA) [1], mean-field DCA (mfDCA) [2] and pseudo-likelihood maximization DCA (plmDCA) [3, 14]. Among that group of methods, mfDCA is very efficient due to the properties of the mean-field approach. The determination of the 2-body couplings consists of a matrix inversion, while the fields are determined through a self-consistency criterion [2]. On the other hand plmDCA usually performs better than mfDCA in terms of accuracy as measured by positive predictive value.

The scores ranking each pair of sites can be conventionally computed using the Frobenius *F*_*ij*_ norm for the couplings between sites *i* and *j*:

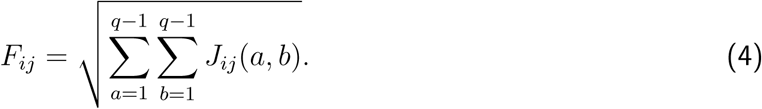

Alternatively, scores can be computed from the direct information *DI_ij_*, which is defined by:

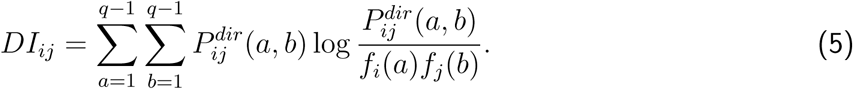

In equation 5, *f*_*i*_(*a*) stands for single-site frequency counts for residue *a* in the MSA and the summation is performed over *q −* 1 states (gap states are excluded). The direct probability 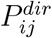 is given as

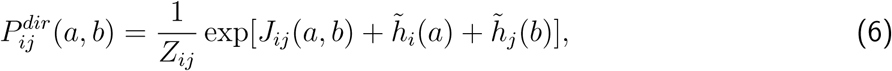

where *Z*_*ij*_ stands for the normalization constant and 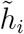 are the single site fields. The DCA scores obtained by using the Frobenius norm or direct information are recomputed using average product correction (APC). Denoting the DCA score by *S*, the average product corrected values are obtained using

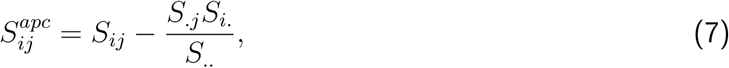

where. denotes the average over the corresponding index.

In mean-field DCA [2], the couplings *J*_*ij*_(*a, b*) are defined as the inverse of the matrix *C*_*ij*_(*a, b*),

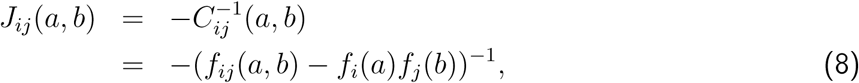

as a function of the single-site *f*_*i*_(*a*) and pair-site *f*_*ij*_(*a, b*) frequencies. These couplings are then used in equation 6 to estimate the two single-site fields (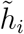 and 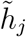) in an iterative procedure by constraining the marginal probabilities of the direct information to be consistent with single-site frequencies as:

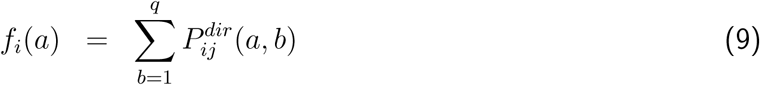

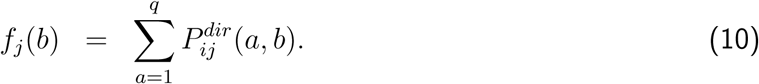

In the pseudolikelihood maximization direct coupling analysis (plmDCA) each site *i* in a sequence *m* is described by a conditional probability of being occupied by a residue 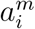 in the presence of residues in other sites as 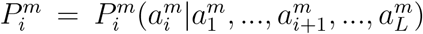. This conditional probability can be written in terms of a new energy function (instead of equation 2) as

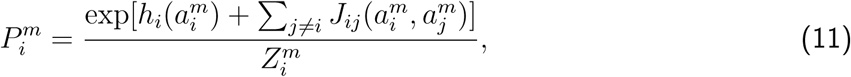

where 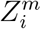 is the normalization constant and is given by

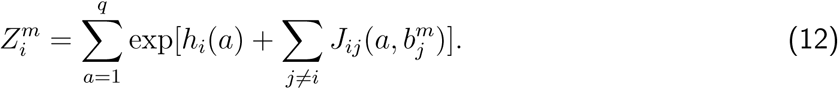

The pseudolikelihood *l* is the product of the conditional probabilities for the entire MSA data. It’s given by

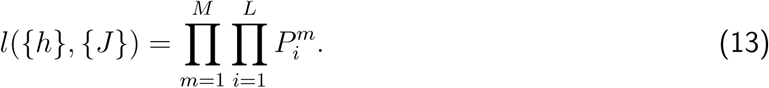

The fields and couplings are estimated by maximizing (or equivalently minimizing the negative) log-pseudolikelihood with regularization. In pydca, we use *L*_2_ norm regularization for both the fields and couplings. The objective function *F* is obtained from the sum of the negative log-pseudolikelihood plus the regularization terms, i.e.,

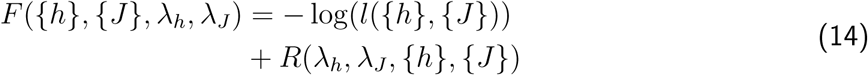

where

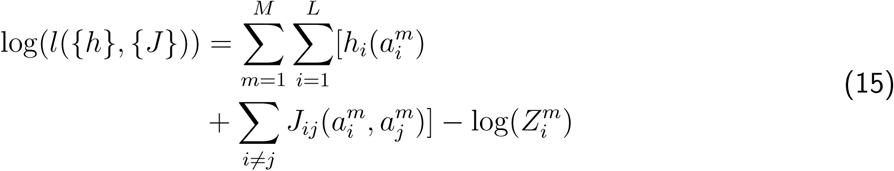

and

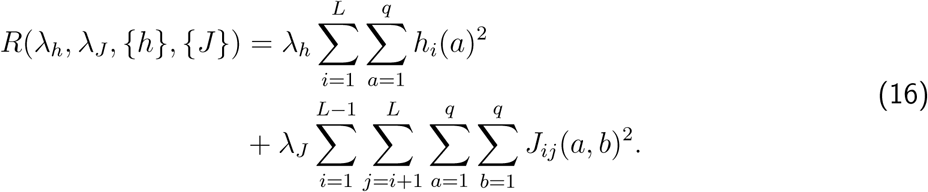

The fields and couplings are estimated by minimizing the objective function *F*. This is done by using gradient descent which requires computation of gradients of the fields and couplings from the objective function. The gradients of *F* are given by

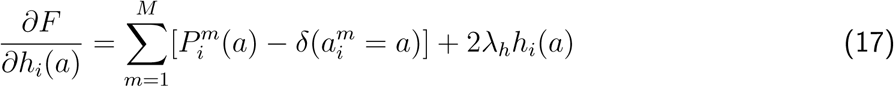

and

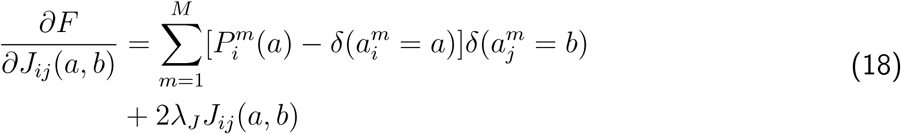

where *δ*(*x* = *y*) is one when *x* = *y* and zero otherwise. The fields and couplings are computed starting from an initial value using iterative procedure. Like the mean-field DCA, scores in pseudolikelihood maximization DCA are quantified either by the Frobenius norm of couplings or by computing the direct information for site-pairs. Note that equations 17 and 18 contain summation over the sequences in the MSA. When sequences are reweighed (see next section) their contribution to the gradients is scaled by the weight. As a result the summation changes from Σ_*m*_ 1 to Σ_*m*_ *ω*_*m*_ where *ω*_*m*_ is the weight of the *m*^*th*^ sequence in the MSA.

The DCA scores deliver quantitative information about the existence of physical contacts in a three-dimensional structure of a biomolecule, where top-scoring pairs are typically used as constraints in molecular modeling tools to predict protein [4, 5, 6, 7, 8, 9, 10, 15, 16, 17, 18] and RNA [11, 12, 19, 20] structures. In addition, the energy function of DCA has been used to study effects of mutations [21, 22, 23, 24]. A summary of the various applications of the DCA method can be found in reference [25].

## Methods: pydca Implementation

### Input Data

The input data for **pydca** is an MSA dataset. **pydca** provides utilities for MSA gap trimming, performs sequence reweighing and a regularization of the dataset.

#### MSA trimming

The accuracy of the DCA computation in general depends on the quality of MSA data. In particular, removing columns in the MSA dataset containing too many gaps reduces the noise, which adversely affects the results. Additionally, **pydca** provides an optional MSA trimming utility, which automatically removes gaps in the MSA columns with respect to the best matching sequence in relation to the reference sequence.

#### Sequence reweighing

From the alignment, single- and double-site frequency counts are computed as

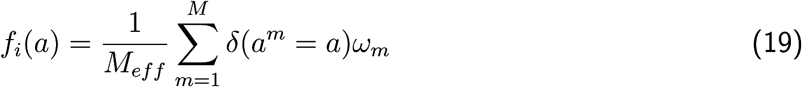

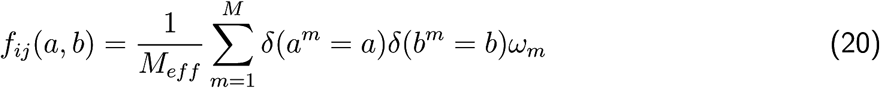

where *a* and *b* denote residue or gap states at sites *i* and *j*, respectively, while *δ*(*x* = *y*) is one when *x* = *y* and zero otherwise. *ω*_*m*_ is the weight of the *m*^*th*^ sequence and *M*_*eff*_ = Σ_*m*_ *ω*_*m*_ is the effective number of sequences in the alignment. The weights of the sequences in the MSA dataset are determined by setting a cut-off value on the sequence identity.

#### Input data regularization

Not all protein or RNA families have a sufficient number of sequences to ensure statistical consistency, such as redundant data or phylogenetic biases. In particular, mean-field DCA matrix inversion may fail due to singularity of the correlation matrix constructed from raw MSA data. To circumvent this problem, input data is regularized by adding pseudocounts to the frequency measures. Using the pseudocounts, the expressions modify to

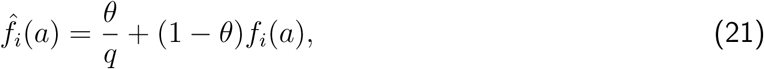

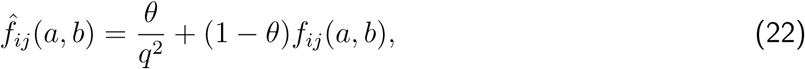

where *θ* controls the pseudocount.

### Algorithm

The two algorithms (mfDCA and plmDCA) in pydca start with the computation of sequence weights, which requires a comparison between individual pairs in the MSA. That formalism contains a time complexity of *O*(*M* ^2^*L*^2^), where *M* is the total number of sequences in the MSA and *L* is the sequence length. The subsequent inversion of the correlation matrix to compute the couplings using mean-field DCA scales with *O*(*L*^3^*q*^3^). If a reference sequence is added, the mapping is done using Biopython’s pairwise sequence alignment utilities [13]. This task has a time complexity of *O*(*ML*_*ref*_ *L*_*min*_), where *L*_*ref*_ is the reference sequence length and *L*_*min*_ is the length of the shortest sequence in the alignment after removal of MSA gaps.

For mean-field DCA, single- and pair-site frequencies are computed from MSA data and stored for the inverse statistical parameter estimation. The highest memory consumption occurs at the storage of the pair-site frequencies and the correlation matrix. These processes consume a storage on the order of *O*(*L*^2^*q*^2^). All other processes require a lower amount of memory.

The plmDCA algorithm requires storage of the fields and couplings, and the corresponding gradients obtained from the objective function. Like the correlation matrix in mean-field DCA, this requires storage of O(*L*^2^*q*^2^). The time complexity scales as O(*L*^4^*q*^3^*MN*) where *N* is the number of gradient descent iterations.

### Usage

#### General Information

As a Python-based package, **pydca** can be used by importing it into other Python source codes or can be installed and used as a command line tool. The command line interface of **pydca** allows to control the input parameters and provides extensive logging messages that help to keep track of the DCA computation progress (see Figure 2). The logging messages are ranked within different levels:

**Figure 1:**
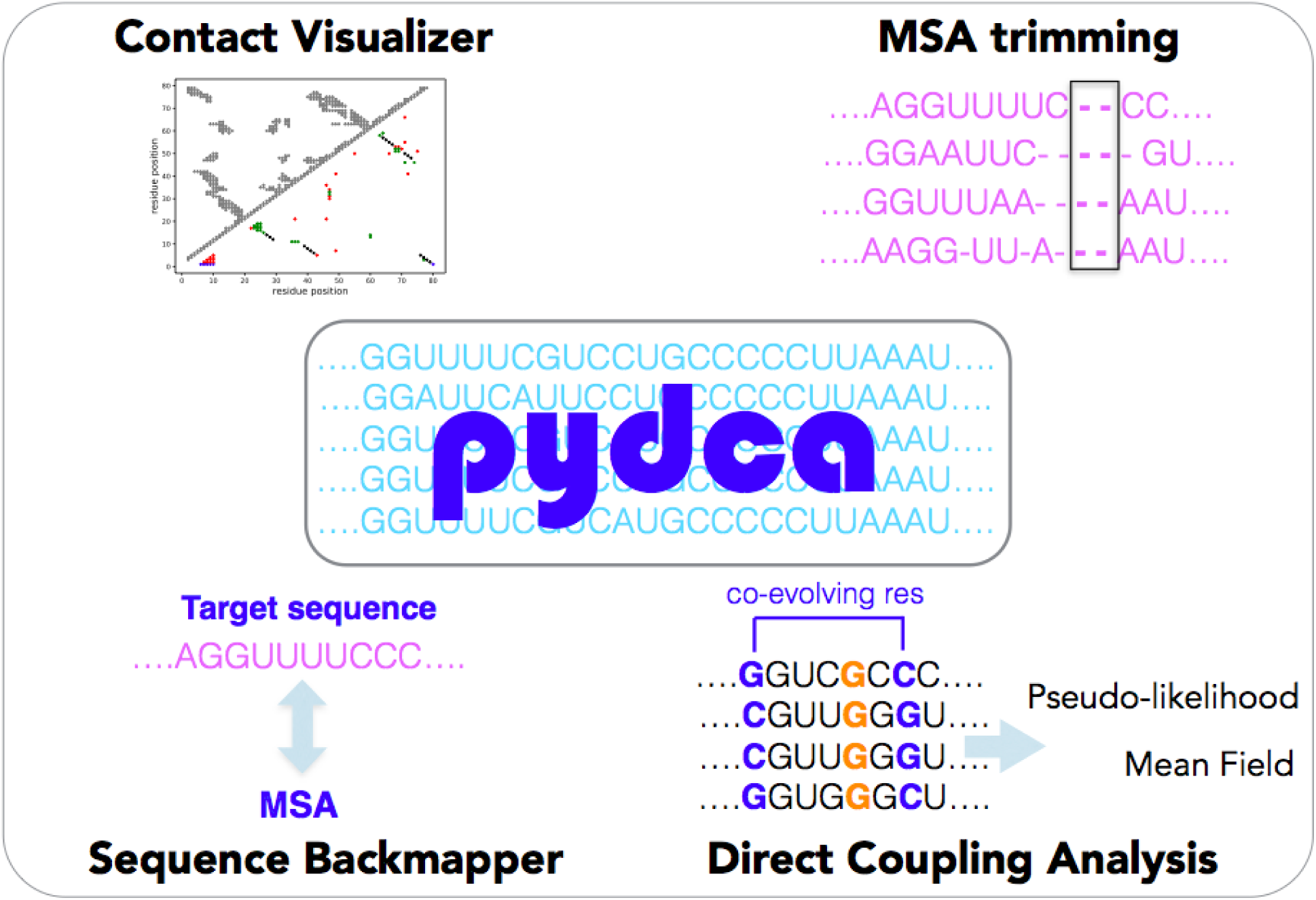
Schematic representation of the main features of the **pydca** software package. The software package combines several functionalities for the contact prediction of proteins and RNA. It includes the sequence backmapping, trimming of the MSA data, inverse inference using the mfDCA and plmDCA algorithms and the final visualization of the DCA result. The software is implemented as a standalone package, simplifying the process of installation and usage.

**Figure 2:**
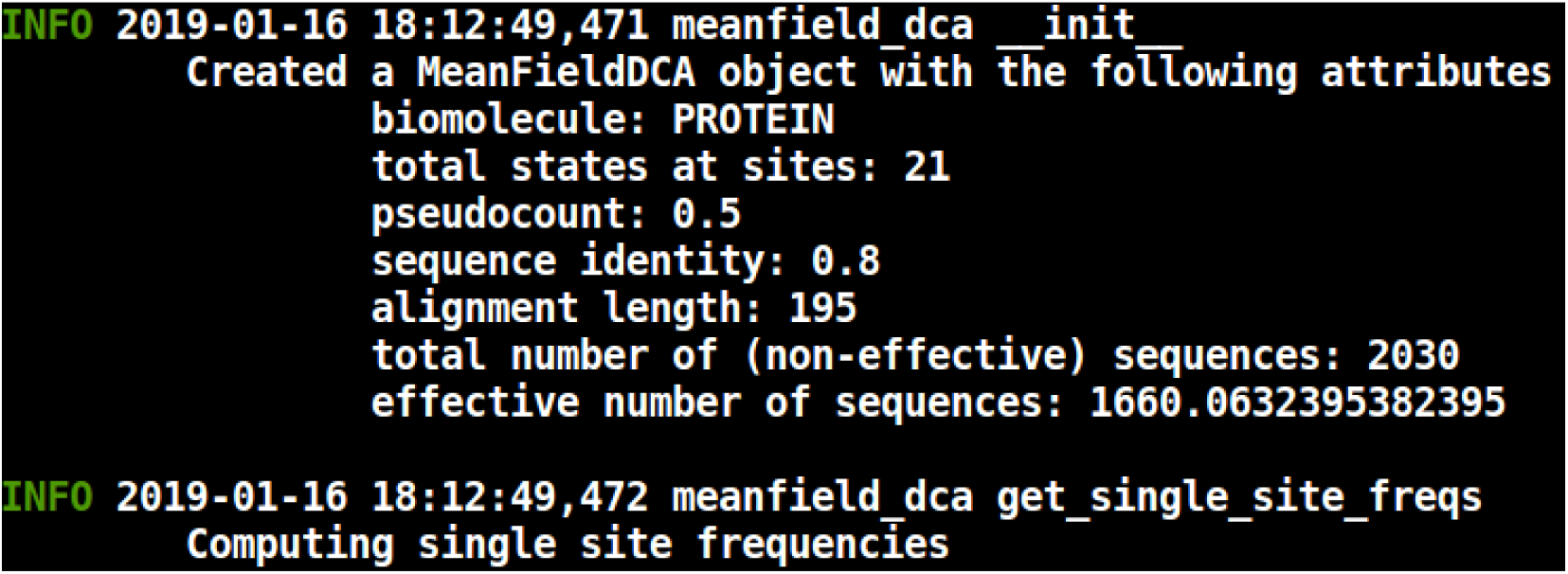
Snapshot of running a DCA computation using the **pydca** software on the command line in verbose mode. Each logging message starts with a logging level (e.g., *INFO*), followed by date and time of display. Afterwards, the name of the module that was used and its corresponding function/method are listed. Task messages are displayed underneath in a new line.

1. *INFO*: These logging messages provide information about DCA computation progress
2. *WARNING*: Logging messages are issued whenever **pydca** encounters ambiguities and resolves them by its own means, e.g., if a reference sequence file contains more than one reference sequence, **pydca** takes the first one and proceeds the computation
3. *ERROR*: Logging messages are signaled when errors occur, e.g., whenever a wrong parameter value is chosen. In case **pydca** detects an error, it sends an indicative error occurred and terminates the computation
4. *MSA trimming command*: Before starting DCA computation, one can curates MSA data by trimming some of the columns of the MSA. For example, all columns in the MSA containing gaps beyond a certain gap threshold are removed:

~~~
pydca trim by gap size <msa file> --max gap 0.9 --verbose
~~~

pydca is one of the main commands provided when **pydca** is installed.

The sub-command `trim by gap size` triggers MSA trimming based on the fraction of gaps found in MSA columns. `<msa_file>` refers to the MSA file supplied by the user. The optional parameter `--max_gap` is used to set the gap threshold (in this case, MSA columns containing more than 90% of gaps will be trimmed) and `--verbose` enables logging messages to be displayed on the terminal (see Figure 2). Another option in the MSA trimming function is defined by:

~~~
pydca trim by refseq <biomolecule> <msa file> <refseq file> --remove all gaps --verbose
~~~

Here, the new subcommand `trim_by_refseq` is used to trim the MSA based on information about the reference sequence obtained from the file `<refseq_file>` supplied by the user. The argument `<biomolecule>` takes either of the values `PROTEIN` or `RNA` (case-insensitive). The optional argument `--remove_all_gaps` removes all gaps in the MSA found in the matching sequence with respect to the reference sequence. If this argument is not supplied, only those gaps in the matching sequence beyond a gap threshold (default 0.5) are removed.

#### DCA computation commands

Coevolutionary scores for pairs of sites in MSA can be obtained from either the Frobenius norm (equation 4) or the direct information (equation 5). To compute the direct information using mean-field DCA, one can execute:

~~~
mfdca compute di <biomolecule> <msa file> --verbose
~~~

The subcommand `compute_di` initiates the computation of DCA scores summarized by direct information. Similarly to compute DCA scores summarized by the Frobenius norm of couplings, we use

~~~
mfdca compute fn <biomolecule> <msa file> --verbose
~~~

The corresponding command for plmDCA is

~~~
plmdca compute fn <biomolecule> <msa file> --num threads 6 --max iterations 500 --verbose
~~~

Here the optional argument `--num_threads` the number of threads for parallel execution. If not set, plmDCA uses a single thread, `--max_iterations` sets the maximum number of iterations for gradient descent.

If a reference sequence file is supplied, the scores corresponding to residue pairs in the reference sequence are computed by mapping the reference sequence to the best matching sequence in the MSA. The best matching sequence in the MSA is searched by pairwise local alignment. When a reference sequence is supplied, the command for mfDCA for computing Frobenius norm is

~~~
mfdca compute fn <biomolecule> <msa file> --refseq file <refseq file> --verbose
~~~

whereas the plmDCA command becomes

~~~
plmdca compute fn <biomolecule> <msa file> --num threads 6 --max iterations 500 --refseq file <refseq file> --verbose
~~~

Furthermore, average product correction can be done for the DCA scores. In **pydca**, this is done by using `--apc` optional argument. For example, using plmDCA

~~~
plmdca compute fn <biomolecule> <msa file> --num threads 4 --max iterations 100 --refseq file <refseq file> --apc --verbose
~~~

#### Visualization command

The result of DCA computation varies based on the input parameters used, e.g., nature of MSA data, regularization and sequence reweighing. The MSA data can be curated in various ways. Regularization is typically done using pseudocounts (equations 21 and 22) for mfDCA or using *L*_2_ norms (equation 16) for plmDCA, whereas sequence reweighing is achieved by setting cut-off value for sequence similarity. To assess how coevolutionary scores are affected by these parameters, **pydca** provides a quick visualization utility. One way of assessing DCA performance is to look up the contact map of a sequence that has an already resolved PDB structure. In **pydca**, this is done on the command line using:

~~~
pydca plot contact map <biomolecule> <pdb chain name> <pdb file> <refseq file> <dca file>
~~~

The new positional argument `<pdb_chain_name>` refers to the chain identifier (chain ID) of the PDB structure chain, `<pdb_file>` the PDB file, `<refseq_file>` the file containing the reference sequence and `<dca_file>` the file containing DCA scores as computed by **pydca**. Note that instead of a PDB file, we can also provide a PDB id and pydca downloads the PDB file from the RCSB PDB database [26]. By default, the top *L* DCA pairs are taken for contact map comparison. These DCA pairs are filtered among all pairs such that they are a few residues apart in the primary sequence. The default separation between these pairs is at least four residues. The user can change this values by using the optional argument `--linear_dist`. Often, PDB files contain multiple chains and it may be difficult to know the existing chains without parsing the PDB file. **pydca** provides a command to display the content of a PDB file when run in verbose mode

~~~
pydca pdb content <pdb file> --verbose
~~~

Another way of visualizing and evaluating DCA scores is to look at the true positive (TP) rate per rank of the predicted pairs.^1^ The TP rate per rank is the number of correctly predicted contacts divided by all predictions at that rank. This can be done in **pydca** by

~~~
pydca plot tp rate <biomolecule> <pdb chain name> <pdb file> <ref file> <dca file>.
~~~

This displays a plot of the TP rate per rank of all ranked pairs. As with the contact map, the default filtering criterion is set to a minimum of four residues in between ranked pairs in the primary sequence. Setting `--linear_dist` to zero overrides this filtering criterion and plots the TP rate of all pairs. Information about all available commands and command line arguments can be accessed via the `--help` optional argument.

#### Use of pydca as library

In addition to the command line interfaces **pydca** can be used as a library by importing it into other python source codes. This is done by importing the desired module of **pydca**. For example, to use the plmDCA algorithm of pydca:

~~~
from pydca.plmdca import plmdca
~~~

Now we have access to the plmdca module, which allows to create a plmDCA object and execute DCA computations in our Python source code. For detailed examples about using **pydca** as a library, we refer to https://github.com/KIT-MBS/pydca

## Results: pydca Application

### Contact Predictions

In this section, we present some examples of DCA computation using the **pydca** software choosing six proteins and six RNAs for validation. The selection criteria of these proteins/RNAs was based on i) existence of sufficient number of sequences for the corresponding families and ii) existence of an experimentally resolved PDB structure so that DCA predicted contacts can be compared with the experimental contact patterns. We considered the following Pfam-sets and PDB structures: PF04542.14 (1or7), PF01497.18 (1n2z), PF00486.28 (2hwv), PF00534.20 (2iv7), PF00691.20 (1oap) and PF13344.6 (2pt1). As RNA families and PDB structures we used RF00059 (2gdi), RF00162 (2gis), RF00167 (1y26), RF00504 (3owi), RF01051 (3irw) and and RF01734 (3vrs).

The sequences of these families were retrieved from the RFAM and PFAM [27, 28] databases and aligned using HMMER (proteins) or infernal (RNA) software [29, 30]. The reference sequences were included in the homologous family if not already present before the alignment. The resulting alignments were trimmed using the reference sequence in a way that only the columns corresponding to reference sequence’s residues were taken into account. All others were discarded as noise.

### Contact map visualization

Here we show the visualization features of **pydca** by a visual comparison of the DCA predicted contact maps with those obtained from the PDB structures for (A) the TPP riboswitch RNA (RF00059, PDB ID 2gdi) and (B) Sigma70 protein (PF04542.14 PDB ID 1or7). For this comparison, we took the top *L* DCA-ranked pairs (see Figure 3).

**Figure 3:**
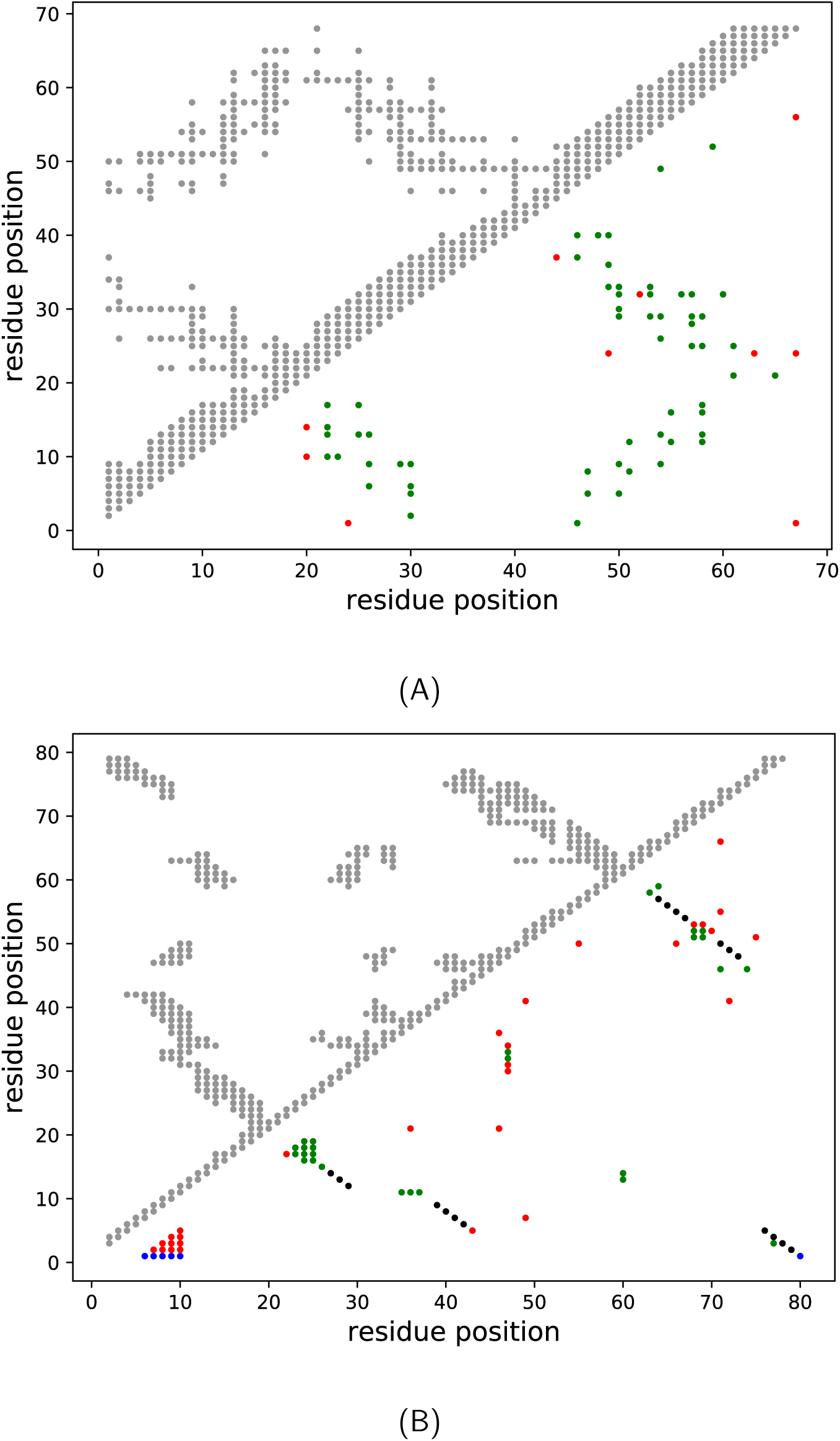
Comparison of **pydca** predicted contacts with PDB contacts. We used the direct-information scores computed using the mean-field algorithm for contact prediction. In (A) we show the PDB ID 1or7 (protein family PF04542.14) and in (B) PDB ID 2gdi (RNA family RF00059). The grey dots are PDB contacts, others are predicted by **pydca**. The red dots are false positives, blue ones are due to missing residues in the PDB structure. The green and black dots are true positives, where the black ones represent the secondary structure pairs in the RFAM consensus secondary structure.

**Figure 4:**
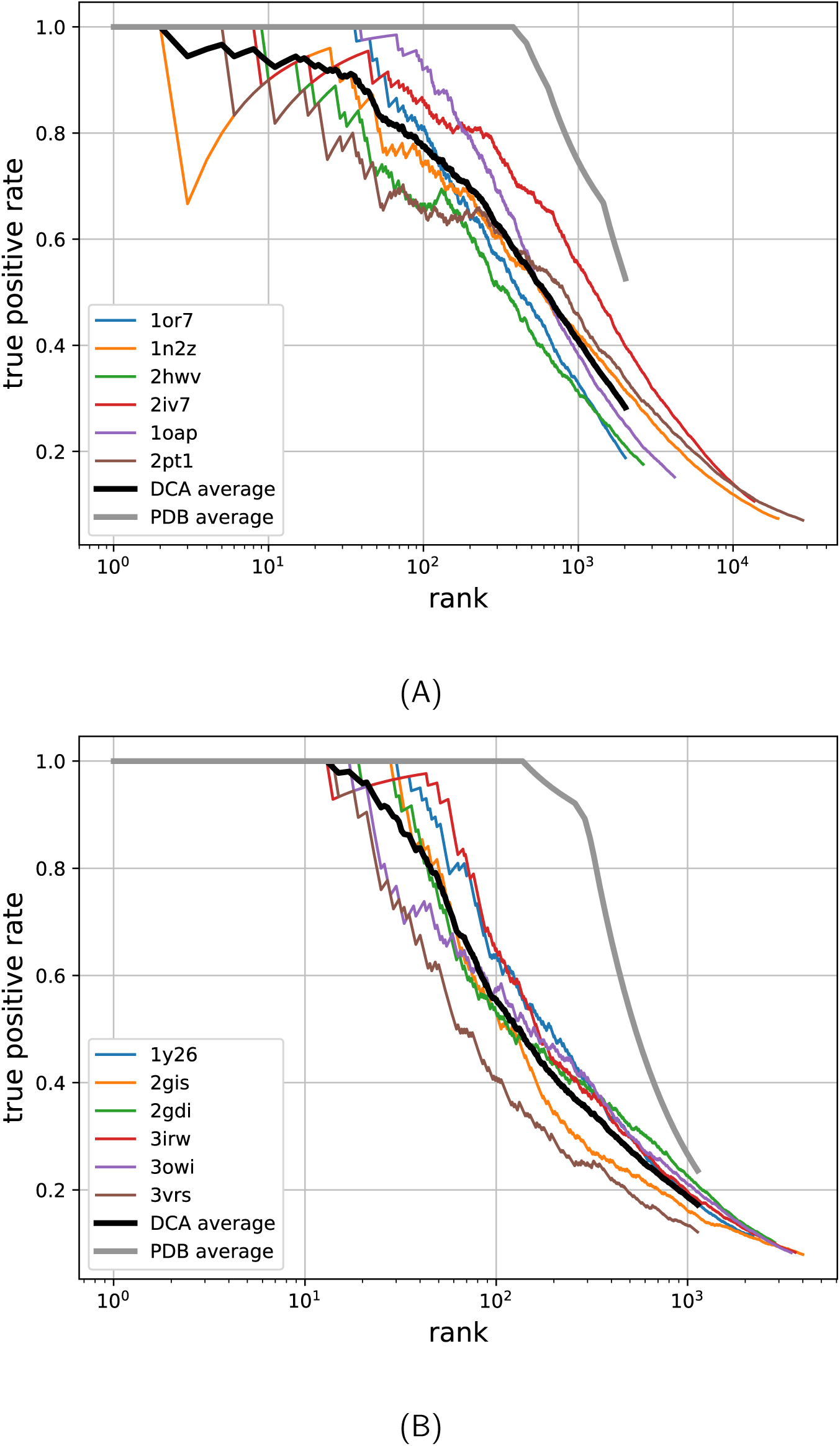
True-positive rates per rank for six protein (A) and six RNA (B) families. The grey line is the average of the true positive rate obtained from PDB structures, representing the theoretically maximum possible true positive rate. The black lines are the average of DCA true positive rates as computed by **pydca**’s mean-field algorithm.

In that figure, the grey dots denote the contact as obtained from PDB structures, others are those of DCA contacts as predicted by **pydca**. The red dots represent false positives and the blue are those missing in PDB that cannot be categorized as false or true positives. The green and black dots are true positives. The black dots represent RNA secondary structure annotations as in the consensus secondary structure of RFAM.

Two residues or two nucleobases are considered to be in physical contact if they have at least a pair of heavy atoms (i.e. non hydrogen atoms) that are less than 8 Å apart in the PDB structure. In addition, we required a minimum sequence distance of 4, i.e. we only considered residue pairs that are not in proximity in the sequence (for sites *i* and *j*, we set |*i* − *j*| > 4).

### Evaluation of pydca performance

Here, we assess the performance of the mean-field and pseudolikelihood maximization algorithms implementation in **pydca**. In addition we compare these two flavor of **pydca** with that of EVcouplings [12, 31] and GREMLIN [32], two pseudo-likelihood maximization DCA methods implemented in C and C++ programming language respectively. We used a Desktop computer with processor Intel corei7-8700 CPU 3.20GHx12 running on Ubuntu 18.04 LTS and setting the number of threads equal to 10.

We performed computation of DCA scores quantified by Frobenius norm with average prodcut correction (APC). The input parameters are set to the values: the sequence identity 0.8, and the pseudocount (value of *θ* in equations 21 and 22) for the mfDCA to be 0.5. The regularization parameters for plmDCA are set to *λ*_*h*_ = 1 and *λ*_*J*_ = 0.2(*L* − 1) for pydca, EVcouplings and GREMLIN. We also set the maximum number of iterations for gradient descent to be large enough so that convergence is reached prior to reaching this limit. Then we computed the true positive rates (TP) of the six proteins and RNAs taking the top *L* DCA ranked site pairs.

Table 1 lists the true positive rates for six protein and RNA families used for comparing the two flavors of pydca implementation (mfDCA and plmDCA), EVcouplings and GREMLIN.

**Table 1:**
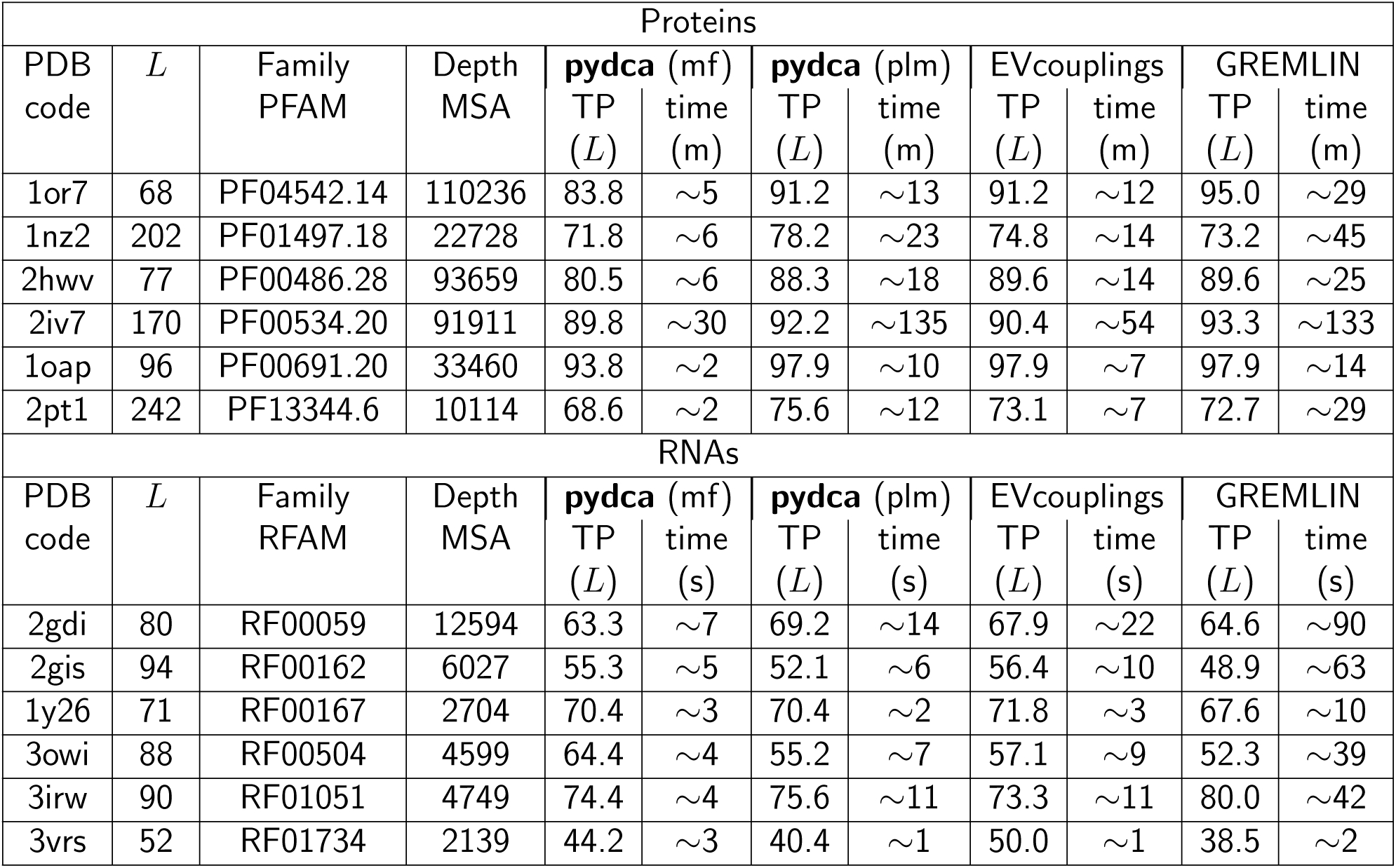
Comparison of method’s performances

Despite the simplicity of the mean-field DCA approach, the computation reaches a very good average TP score of about 80% and 60% for the top-*L* contacts considered in the protein and RNA structures respectively. Moreover, the computation is also very fast running all these middle-size entries in about 50 minutes for proteins or less then half-minute for RNAs, making mfpydca the perfect tool for analyzing large amount of data.

The three pseudo-likelihood implementations are slightly more accurate than the mean-field DCA approach, in particular for proteins where there is a statistical significant improvement of the TP rate score of about 7%. No significant difference between mean-field and the pseudo-likelihood approaches has been observed for the six RNAs. Our implementation (plmpydca) has on average the same accuracy of EVcouplings and GREMLIN. Speed-wise, plmpydca is slightly slower than EVcoupling for proteins and slightly faster for RNAs. plmpydca outperforms GREMLIN in terms of speed for both proteins and RNAs.

## Acknowledgments

We thank the *Impuls- und Vernetzungfond* of the Helmholtz Association. We also thank Martin Weigt for useful discussions.

## Footnotes

1 Naturally, this analysis can only be done if the real contacts are known, e.g. from a biomolecular structure. The TP rate therefore merely tests the quality of contact prediction.

## References

[1] Martin Weigt, Robert a White, Hendrik Szurmant, James a Hoch, and Terence Hwa. Identification of direct residue contacts in protein-protein interaction by message passing. Proc. Natl. Acad. Sci. U.S.A., 106(1):67–72, 2009.

[2] Faruck Morcos, Andrea Pagnani, Bryan Lunt, Arianna Bertolino, Debora S Marks, Chris Sander, Riccardo Zecchina, José N Onuchic, Terence Hwa, and Martin Weigt. Direct-coupling analysis of residue coevolution captures native contacts across many protein families. Proc. Natl. Acad. Sci. U.S.A., 108(49):E1293–301, 2011.

[3] Magnus Ekeberg, Cecilia Lövkvist, Yueheng Lan, Martin Weigt, and Erik Aurell. Improved contact prediction in proteins: Using pseudolikelihoods to infer Potts models. Phys. Rev. E - Stat. Nonlin. Soft Mat. Phys., 87(1):1–16, 2013.

[4] Ricardo N. dos Santos, Faruck Morcos, Biman Jana, Adriano D. Andricopulo, and José N. Onuchic. Dimeric interactions and complex formation using direct coevolutionary couplings. Sci. Rep., 5(1):13652, 2015.

[5] Alexander Schug, Martin Weigt, José N Onuchic, Terence Hwa, and Hendrik Szurmant. High-resolution protein complexes from integrating genomic information with molecular simulation. Proc. Natl. Acad. Sci. U.S.A., 106(52):22124–22129, 2009.

[6] Debora S Marks, Lucy J Colwell, Robert Sheridan, Thomas A Hopf, Andrea Pagnani, Riccardo Zecchina, and Chris Sander. Protein 3D structure computed from evolutionary sequence variation. PLoS ONE, 6(12), 2011.

[7] Joanna I Sułkowska, Faruck Morcos, Martin Weigt, Terence Hwa, and José N Onuchic. Genomics-aided structure prediction. Proc. Natl. Acad. Sci. U.S.A., 109(26):10340–10345, 2012.

[8] Thomas A Hopf, Lucy J Colwell, Robert Sheridan, Burkhard Rost, Chris Sander, and Debora S Marks. Three-dimensional structures of membrane proteins from genomic sequencing. Cell, 149(7):1607–1621, 2012.

[9] Angel E Dago, Alexander Schug, Andrea Procaccini, James a Hoch, Martin Weigt, and Hendrik Szurmant. Structural basis of histidine kinase autophosphorylation deduced by integrating genomics, molecular dynamics, and mutagenesis. Proc. Natl. Acad. Sci. U.S.A., 109(26):E1733–42, 2012.

[10] F. Morcos, B. Jana, T. Hwa, and J. N. Onuchic. Coevolutionary signals across protein lineages help capture multiple protein conformations. Proc. Natl. Acad. Sci. U.S.A., 110(51):20533–20538, 2013.

[11] Eleonora De Leonardis, Benjamin Lutz, Sebastian Ratz, Simona Cocco, Rémi Monasson, Alexander Schug, and Martin Weigt. Direct-Coupling Analysis of nucleotide coevolution facilitates RNA secondary and tertiary structure prediction. Nucl. Acids Res., 43(21):10444–10455, 2015.

[12] Caleb Weinreb, Adam J. Riesselman, John B. Ingraham, Torsten Gross, Chris Sander, and Debora S. Marks. 3D RNA and Functional Interactions from Evolutionary Couplings. Cell, 165(4):963–975, 2016.

[13] Peter J. A. Cock, Tiago Antao, Jeffrey T. Chang, Brad A. Chapman, Cymon J. Cox, Andrew Dalke, Iddo Friedberg, Thomas Hamelryck, Frank Kauff, Bartek Wilczynski, and Michiel J. L. de Hoon. Biopython: freely available python tools for computational molecular biology and bioinformatics. Bioinformatics, 25(11):1422–1423, 2009.

[14] Erik Aurell and Magnus Ekeberg. Inverse ising inference using all the data. Phys. Rev. Lett., 108(9):1–5, 2012.

[15] Agnes Toth-Petroczy, Perry Palmedo, John Ingraham, Thomas A Hopf, Bonnie Berger, Chris Sander, and Debora S Marks. Structured states of disordered proteins from genomic sequences. Cell, 167(1):158–170, 2016.

[16] Yuefeng Tang, Yuanpeng Janet Huang, Thomas A Hopf, Chris Sander, Debora S Marks, and Gaetano T Montelione. Protein structure determination by combining sparse NMR data with evolutionary couplings. Nat. Meth., 12(8):751–754, 2015.

[17] Sikander Hayat, Chris Sander, Debora S Marks, and Arne Elofsson. All-atom 3d structure prediction of transmembrane β-barrel proteins from sequences. Proc. Natl. Acad. Sci. U.S.A., 112(17):5413–5418, 2015.

[18] Guido Uguzzoni, Shalini John Lovis, Francesco Oteri, Alexander Schug, Hendrik Szurmant, and Martin Weigt. Large-scale identification of coevolution signals across homo-oligomeric protein interfaces by direct coupling analysis. Proc. Natl. Acad. Sci. U.S.A., 114(13):E2662–E2671, 2017.

[19] Jian Wang, Kangkun Mao, Yunjie Zhao, Chen Zeng, Jianjin Xiang, and Yi Zhang. Optimization of RNA 3D structure prediction using evolutionary restraints of nucleotide-nucleotide interactions from direct coupling analysis. Nucl. Acids Res., 2017.

[20] Fabrizio Pucci and Alexander Schug. Shedding light on the dark matter of the biomolecular structural universe: Progress in RNA 3D structure prediction. Methods, 2019.

[21] Matteo Figliuzzi, Herve Jacquier, Alexander Schug, Oliver Tenaillon, and Martin Weigt. Coevolutionary landscape inference and the context-dependence of mutations in beta-lactamase tem-1. Mol. Biol. Evol., 33(1):268–280, 2016.

[22] Ryan R Cheng, Olle Nordesjö, Ryan L Hayes, Herbert Levine, Samuel C Flores, José N Onuchic, and Faruck Morcos. Connecting the sequence-space of bacterial signaling proteins to phenotypes using coevolutionary landscapes. Mol. Biol. Evol., 33(12):3054–3064, 2016.

[23] William F Flynn, Allan Haldane, Bruce E Torbett, and Ronald M Levy. Inference of epistatic effects leading to entrenchment and drug resistance in hiv-1 protease. Mol. Biol. Evol., pages 10–1093, 2017.

[24] Thomas A Hopf, John B Ingraham, Frank J Poelwijk, Charlotta PI Schärfe, Michael Springer, Chris Sander, and Debora S Marks. Mutation effects predicted from sequence co-variation. Nat. Biotechnol., 35(2):128–135, 2017.

[25] Mehari B. Zerihun and Alexander Schug. Biomolecular coevolution and its applications: Going from structure prediction toward signaling, epistasis, and function. Biochem. Soc. Trans., 45(6):1253–1261, 2017.

[26] Helen M. Berman, John Westbrook, Zukang Feng, Gary Gilliland, T. N. Bhat, Helge Weissig, Ilya N. Shindyalov, and Philip E. Bourne. The Protein Data Bank. Nucleic Acids Research, 28(1):235–242, 01 2000.

[27] Ioanna Kalvari, Joanna Argasinska, Natalia Quinones-Olvera, Eric P Nawrocki, Elena Rivas, Sean R Eddy, Alex Bateman, Robert D Finn, and Anton I Petrov. Rfam 13.0: shifting to a genome-centric resource for non-coding rna families. Nucl. Acids Res., 46(D1):D335–D342, 2018.

[28] Robert D. Finn, Penelope Coggill, Ruth Y. Eberhardt, Sean R. Eddy, Jaina Mistry, Alex L. Mitchell, Simon C. Potter, Marco Punta, Matloob Qureshi, Amaia Sangrador-Vegas, Gustavo A. Salazar, John Tate, and Alex Bateman. The Pfam protein families database: Towards a more sustainable future. Nucl. Acids Res., 44(D1):D279–D285, 2016.

[29] Robert D. Finn, Jody Clements, and Sean R. Eddy. HMMER web server: interactive sequence similarity searching. Nucl. Acids Res., 39(suppl2):W29–W37, 05 2011.

[30] Eric P. Nawrocki and Sean R. Eddy. Infernal 1.1: 100-fold faster RNA homology searches. Bioinformatics, 29(22):2933–2935, 09 2013.

[31] Thomas A Hopf, Anna G Green, Benjamin Schubert, Sophia Mersmann, Charlotta P I Schärfe, John B Ingraham, Agnes Toth-Petroczy, Kelly Brock, Adam J Riesselman, Perry Palmedo, Chan Kang, Robert Sheridan, Eli J Draizen, Christian Dallago, Chris Sander, and Debora S Marks. The EVcouplings Python framework for coevolutionary sequence analysis. Bioinformatics, 35(9):1582–1584, 10 2018.

[32] Hetunandan Kamisetty, Sergey Ovchinnikov, and David Baker. Assessing the utility of coevolution-based residue–residue contact predictions in a sequence-and structure-rich era. Proceedings of the National Academy of Sciences, 110(39):15674–15679, 2013.

